# Developmental Venous Anomalies are a Genetic Primer for Cerebral Cavernous Malformations

**DOI:** 10.1101/2021.08.25.457657

**Authors:** Daniel A. Snellings, Romuald Girard, Rhonda Lightle, Abhinav Srinath, Sharbel Romanos, Ying Li, Chang Chen, Aileen A. Ren, Mark L. Kahn, Issam A. Awad, Douglas A. Marchuk

## Abstract

Cerebral cavernous malformations (CCM) are a neurovascular anomaly that may occur sporadically in otherwise healthy individuals, or be inherited by autosomal dominant mutations in the genes that encode the proteins of the CCM signaling complex (*KRIT1, CCM2*, or *PDCD10*)^1–4^. CCMs have long been known to follow a genetic two-hit model where lesion formation is initiated by somatic mutations resulting in biallelic loss of a CCM complex gene^5–8^. Recent studies have shown that somatic mutations in *MAP3K3* and *PIK3CA* also contribute to CCM pathogenesis^9–11^; however, it remains unclear how these mutations contribute to sporadic versus familial cases. Here we show that somatic mutations in *MAP3K3* are mutually exclusive with mutations in CCM complex genes and that mutations in *MAP3K3* contribute to sporadic, but not familial CCM. Using single-nucleus DNA sequencing, we show that co-occurring *MAP3K3* and *PIK3CA* mutations are present within the same clonal population of cells. Furthermore, we identify *PIK3CA* mutations in CCM-associated developmental venous anomalies (DVA). It has long been known that sporadic CCM often develop in the vicinity of a DVA. However, the underlying cause of this association is unknown^12–14^. In this first report of the molecular pathology of CCM-associated DVA, we find that the identical *PIKC3A* mutation is found in both the DVA and its associated CCM, but that an activating *MAP3K3* mutation appears only in the CCM. These results support a mechanism where DVA develop as the result of a *PIK3CA* mutation, creating a region of the brain vasculature that functions as a genetic primer for CCM development following acquisition of an additional somatic mutation.

## Introduction

Cerebral cavernous malformations (CCMs) are hemorrhagic neurovascular malformations that may lead to stroke, seizures and other clinical sequelae. CCM disease may be inherited by an autosomal dominant loss of function (LOF) mutation in the genes encoding components of the CCM signaling complex: *KRIT1*^2,4^, *CCM2* ^3^, or *PDCD10* ^1^. Solitary sporadic CCM lesions may also occur in the absence of inherited germline mutations in CCM complex genes. Previous studies have established that CCM pathogenesis follows a genetic two-hit model where somatic mutations cause biallelic LOF in *KRIT1, CCM2*, or *PDCD10* to initiate lesion formation^5–8^. Recent studies have found that somatic mutations in *PIK3CA* and *MAP3K3* also contribute to CCM pathogenesis^9–11^. These discoveries opened new avenues for research and therapeutic development, but they also raised new questions about the roles of somatic mutations in familial versus sporadic lesions.

The presence of multiple somatic mutations in CCMs also helps explain a long-standing mystery: the association between sporadic CCM lesions and developmental venous anomalies (DVA). DVA are the most common vascular malformation present in 6-14% of the adult population^15–18^ with the majority developing prior to the age of 20^15^. When assessed by magnetic resonance imaging, an adjacent DVA is identified in 24-32% of sporadic CCM cases^12–14^, and an even greater fraction of sporadic CCMs are found to be associated with a DVA at surgery^12,13^. One study focused on DVA reported an adjacent sporadic CCM in 6.9% of all DVAs in a general population (116 of 1689)^15^. These studies highlight the association between DVA and sporadic CCM. By contrast, familial CCM lesions have not been associated with DVA^19^. These combined data suggest that a DVA is not required for CCM formation but may be a predisposing factor in sporadic cases. We hypothesize that DVA are caused by somatic mutations, and that the genetic origin of DVA overlaps that of CCM. In this study we probe the relationship between mutations in *KRIT1, CCM2, PDCD10, PIK3CA*, and *MAP3K3* and explore a mechanism by which DVA act as a genetic primer for the genesis of sporadic CCM lesions.

## Results

### Mutations in MAP3K3, KRIT1, CCM2, and PDCD10, are mutually exclusive

To evaluate whether sporadic and familial CCMs have distinct somatic mutation spectra we identified somatic mutations present in 71 CCMs (20 familial CCMs and 51 sporadic/presumed sporadic CCMs). Mutations in *KRIT1, CCM2, PDCD10*, and *PIK3CA* were detected by targeted sequencing and/or droplet digital PCR (ddPCR) as previously described^10^. The common gain of function mutation in *MAP3K3* (hg38 chr17:63691212, NM_002401.3, c.1323C>G; NP 002392, p.I441M) was detected by ddPCR using a previously published probe set^20^.

The p.I441M mutation in *MAP3K3* was identified in 15/51 sporadic CCMs and 0/20 familial CCMs (Figure 1A). We also screened for *MAP3K3* p.I441M in 8 blood samples for which we were previously unable to identify a germline mutation in *KRIT1, CCM2, PDCD10*. None of the 8 blood samples harbored *MAP3K3* p.I441M. Notably 11/51 sporadic CCMs harbored at least 1 somatic mutation in *KRIT1, CCM2*, or *PDCD10*, however none of these CCMs also had a mutation in *MAP3K3* indicating that a mutual loss of the CCM complex and gain of function in MEKK3 (the protein product of *MAP3K3*) are not both required for CCM formation. As the CCM complex is known to be a direct inhibitor of MEKK3 activity ^21,22^, these data strongly suggest identical functional consequences of these mutations.

**Figure 1:**
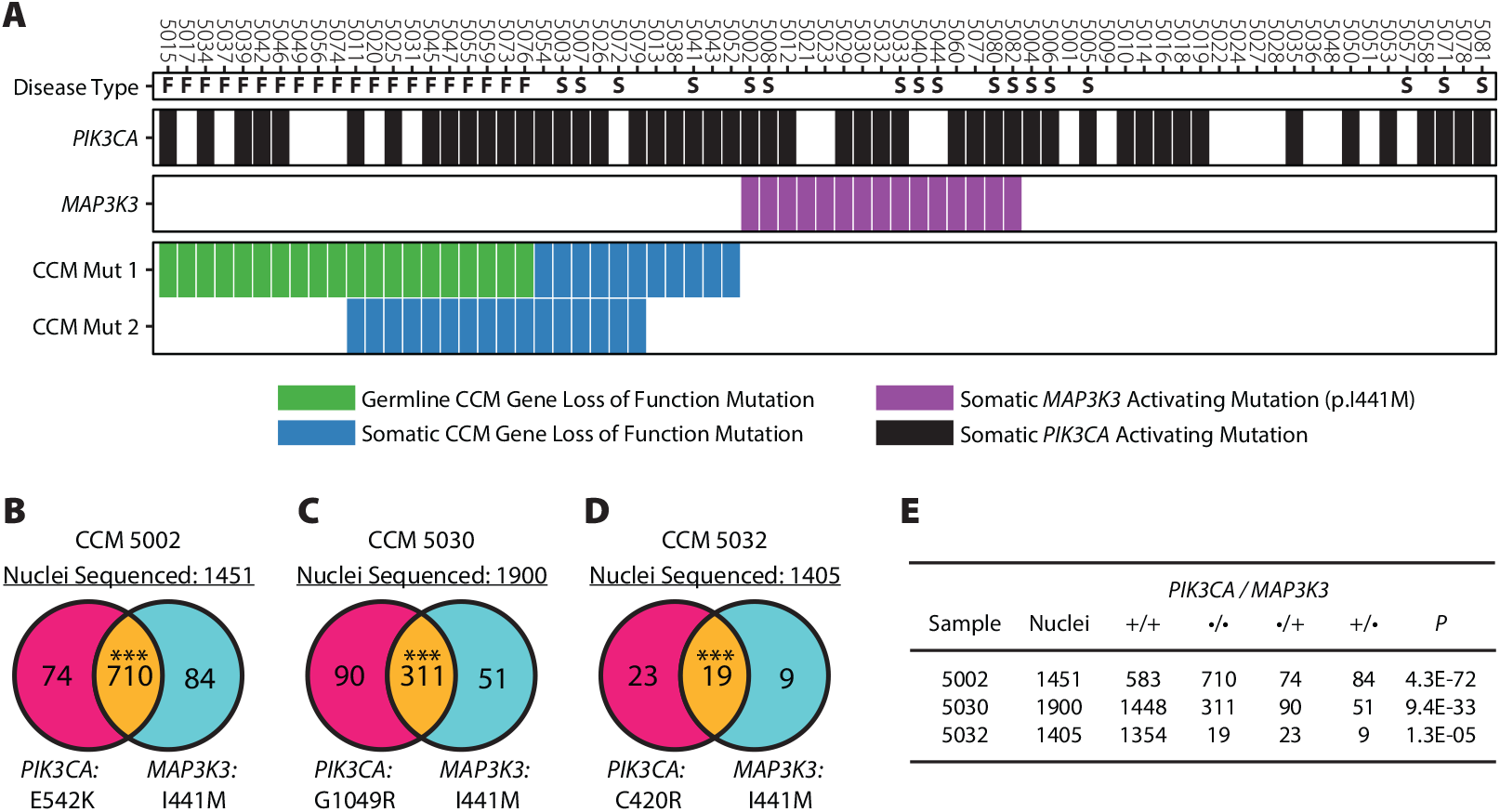
Mutations in *MAP3K3* are mutually exclusive with CCM mutations and occur in the same cells as *PIK3CA* mutations. **A**. Mutations present in 71 CCM samples. Disease type denotes whether the sample was familial (F), sporadic (S), or unknown (blank). The presence of somatic mutations in *PIK3CA* and *MAP3K3* are denoted by black and purple bars respectively. Germline and somatic mutations (green and blue respectively) in *KRIT1, CCM2*, or *PDCD10*, are shown in CCM Mut 1 with the second-hit mutation shown in CCM Mut 2 if present. **B-D**. Nuclei genotypes determined by snDNA-seq. The left and right circles in each Venn diagram shows the number of nuclei with the *PIK3CA* or *MAP3K3* mutations where the overlap shows nuclei harboring both mutations. *** P < 0.0001. **E**. Summary of data presented in B-D including P values determined by comparing the observed number of double mutant nuclei to the expected value derived from a Poisson distribution as done previously ^10^.

The majority of CCM and verrucous venous malformations with a mutation in *MAP3K3* harbor the p.I441M variant^9,11,20^, however an alternative variant p.Y544H has also been identified in a venous malformation^23^. While ddPCR provides superior sensitivity and specificity compared to targeted sequencing, it is restricted to detecting a single mutation per assay. To determine whether other mutations that contribute to CCM pathogenesis—either *MAP3K3* mutations besides p.I441M, or mutations in yet undiscovered genes—we performed whole-exome sequencing (mean depth 133x) on 8 sporadic CCMs for which no somatic mutations in *KRIT1, CCM2, PDCD10*, or *MAP3K3* were found. No additional mutations in *MAP3K3* were identified and no candidate variants in other genes passed QC filters (see Methods).

### Somatic mutations in MAP3K3 and PIK3CA occur in the same population of cells

While somatic mutations in *KRIT1, CCM2, PDCD10*, and *MAP3K3* are mutually exclusive, somatic gain of function mutations in *PIK3CA* may co-occur with any other mutation (Figure 1A). We have previously shown that co-occurring mutations in *KRIT1*/*CCM2* and *PIK3CA* occur in the same clonal population of cells^10^. To determine whether *MAP3K3* and *PIK3CA* mutations co-exist in the same cells we performed single-nucleus DNA-sequencing (snDNA-seq) on frozen tissue from three surgically resected CCMs determined to harbor both mutations (Figure 1B-D).

In CCMs 5002 and 5030, the vast majority of mutant nuclei harbor both mutations in *MAP3K3* and in *PIK3CA* indicating that these mutations co-exist in the same cells. In CCM 5032, 37% (19/51) of mutant nuclei harbor both mutations. While this is a far lower fraction compared to other samples, it is significantly higher than may be expected by chance when sampling from 1405 total nuclei (P = 1.3E-05, Figure 1E). In bulk genetic analysis, the allele frequencies of *PIK3CA* and *MAP3K3* mutations detected in CCM 5002 were 19% and 13% respectively. In snDNA-seq the allele frequencies of these mutations increased to 54% and 55% respectively. This difference likely reflects the mosaic nature of CCMs. As snDNA-seq requires nuclei harvested from frozen tissue, we must sample a new area of the frozen lesion than was sampled for bulk sequencing. Sampling from different sites of the same lesion often results in minor changes in allele frequency, however the drastic change in allele frequency we find in CCM 5002 suggests either that our initial sample of the lesion for bulk sequencing contained largely non-lesion tissue, or an uneven distribution of mutant cells in the lesion.

### CCMs and associated Developmental Venous Anomalies harbor a shared mutation in PIK3CA

Many sporadic CCMs are found in the vicinity of a developmental venous anomaly (DVA). We hypothesized that CCM and DVA may have a common genetic origin, specifically that DVA may be a genetic precursor to CCM. To determine whether DVA and CCMs originate from a shared mutation, we collected three sporadic CCMs and sampled a portion of the associated DVA obtained during surgery (Figure 2A-C). Assays for mutations via ddPCR revealed that all three CCMs have a somatic activating mutation in *PIK3CA* and that the same mutation is present within the paired DVA samples at lower frequency (Figure 2D). Furthermore, ddPCR revealed that two of the CCMs harbored a mutation in *MAP3K3* in addition to the previously noted mutation in *PIK3CA*. However, unlike the *PIK3CA* mutation, the *MAP3K3* mutation was entirely absent from both DVA samples (Figure 2E). The presence of the *PIK3CA*, but not the *MAP3K3*, mutation in the DVA confirms that the *PIK3CA* mutation in the DVA did not arise via cross-sample contamination. The presence of multiple somatic mutations in these CCMs allows us to infer the developmental history of the lesion. The cancer field commonly uses the presence or absence of somatic mutations in clonal populations to track the evolutionary history of a tumor^24,25^. Recent studies have expanded on this approach to use somatic mutations as endogenous barcodes to track embryonic development^26^. Using this same approach, we infer that the DVA was the first lesion to develop and that the associated CCM is derived from cells of the DVA following a somatic mutation in *MAP3K3*.

**Figure 2:**
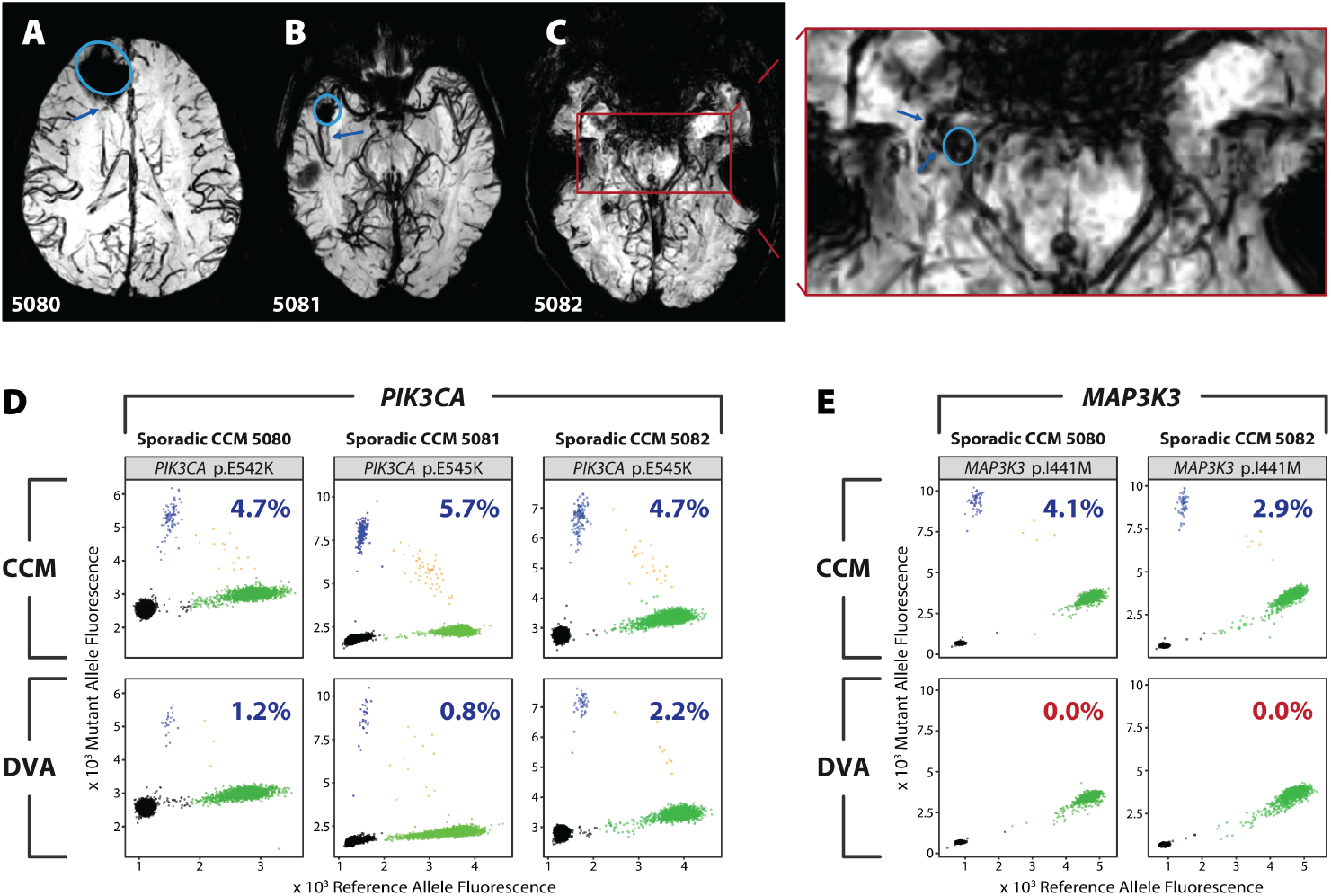
Associated CCM and DVA harbor identical somatic mutations in *PIK3CA*. **A, B, C**. Axial magnetic resonance (MR) susceptibility weighted images acquired at 3 Tesla showing CCM (circle) and associated DVA branches sampled during surgery (arrow) in individuals with CCM 5080 (A) or 5081 (B) or 5082(C). The inset red box in C shows the region expanded to the right with the CCM and DVA marked. **D, E**. Somatic mutations in *PIK3CA* (D) and *MAP3K3* (E) in CCM (top panels) and the associated DVA (bottom panels) from samples 5081, 5082, and 5083. Mutations were detected by droplet digital PCR (ddPCR) and shown as the fluorescence of the reference probe on the x-axis, and the mutant probe on the y-axis. Droplets containing the reference allele, mutant allele, both, or neither, are colored in green, blue, orange, and black respectively. Percentage inset into each graph shows the variant allele frequency for the displayed mutation. If the mutation was determined to be present, the percentage is blue, else the percentage is red.

### Individuals with DVAs alone have differentially expressed (DE) plasma miRNAs related to PIK3CA and -B, while individuals with sporadic CCM and associated DVA have DE plasma miRNAs related to PIK3CA, AKT3, and MAP3K3

In addition to assaying the presence of *PIK3CA* mutations in DVA associated with CCM, we would ideally also assay DVA that are not associated with CCM. Unfortunately, DVA are benign malformations and are not resected unless associated with an additional pathology. This has precluded the direct assessment of *PIK3CA* mutations in DVA without a CCM. To address this limitation, we sought another source of tissue that could be assayed for indirect evidence of *PIK3CA* activation. Thus, we collected plasma from individuals with DVA without a CCM and measured circulating miRNAs that might serve as biomarkers reflecting *PIK3CA* activity^27^.

We sequenced the plasma miRNomes of 12 individuals with a sporadic CCM associated with a DVA (CCM + DVA), 6 individuals with a DVA without a CCM (DVA only), and 7 healthy controls. Three plasma miRNAs were DE in the DVA only group when compared to healthy controls (P < 0.05; false discovery rate (FDR) corrected). Two of these DE miRNAs, *miR-134-5p* (FC = 0.10) and *miR-92a-3p* (FC = 3.10), putatively target *PIK3CA* and *PIK3CB* respectively, which are both components of the PI3K/AKT/mTOR pathway (Supplemental Figure 1).

In addition, 18 plasma miRNAs were DE in CCM + DVA when compared to DVA only (P < 0.05; FDR corrected). Two of the 18 DE miRNAs, *miR-122-5p* (FC = 5.25) and *miR-182-5p* (FC = 0.21), target *AKT3* and *MAP3K3* respectively, linking them to both the PI3K/AKT/mTOR and MAPK/ERK pathways. Additionally, *let-7c-5p* (FC = 12.64) targets both *PIK3CA* and *MAP3K3*, consistent with the role of these two pathways in sporadic CCM pathogenesis (Supplemental Figure 1). Of interest, *let-7c-5p* also targets *COL1A1*, a DEG within the transcriptome of human sporadic CCM lesions (see Supplementary Data). This gene is associated with PI3K/AKT signaling, platelet activation, and ECM-receptor interaction KEGG pathways^28,29^, all of which have previously been implicated in CCM disease^9,10,30,31^.

Additionally, 28 DE plasma miRNAs were identified between CCM + DVA and healthy controls (P < 0.05; FDR corrected). Four of these miRNAs putatively target *PIK3CA*: *miR-148a-3p* (FC = 3.27), *miR-148b-3p* (FC = 2.64), *miR-128-3p* (FC = 2.55) and *let-7c-5p* (FC = 4.20), which also targets *MAP3K3*.

### Relative probability of mutations in KRIT1, CCM2, PDCD10, MAP3K3, and PIK3CA

The association between DVA and sporadic—but not familial—CCMs suggests that there are at least two genetic trajectories by which a CCM may develop. The first trajectory is via a quiescent CCM caused by an initial mutation in *KRIT1, CCM2, PDCD10*, or *MAP3K3*. The second trajectory is via a DVA caused by an initial mutation in *PIK3CA* with subsequent mutations in *KRIT1, CCM2, PDCD10*, or *MAP3K3* leading to CCM formation. To understand whether one trajectory is favored in familial vs sporadic CCMs we use a simplified model to estimate the relative probability of CCM complex LOF (*KRIT1, CCM2, PDCD10*), *MAP3K3* GOF, and *PIK3CA* GOF.

Somatic mutagenesis is a complex process that depends on many variables including cell turnover rates, age, exposure to mutagens, and numerous genetic factors which facilitate DNA synthesis and repair. As a result, estimating absolute mutation rate is not feasible. However, by assuming a uniform mutation rate across these genes, we can determine the relative rate of CCM complex LOF, *MAP3K3* GOF, and *PIK3CA* GOF as simply the number of mutations that result in LOF or GOF. Determining this value for *MAP3K3* and *PIK3CA* is straightforward. To date there is only a single mutation in *MAP3K3* (p.I441M) that has been reported in CCMs. The spectrum of mutations in *PIK3CA* has been well documented in the catalogue of somatic mutations in cancer (COSMIC). In the absence of functional assays for each mutation, we define a GOF mutation as any mutation that accounts for >1% of all *PIK3CA* mutations COSMIC which was determined to be 10 mutations. The number of CCM complex LOF mutations was determined by considering all possible single nucleotide variants that would result in a premature stop codon or disrupt a canonical splice site (see Methods). The average of these values for all three genes is 430 mutations (*KRIT1* = 718, *CCM2* = 353, *PDCD10* = 220). This is a very conservative estimate and should be considered a lower bound, since the majority of CCM complex LOF mutations are the result of frameshift mutations which are not accounted for here.

From the relative probability of each mutation, it is clear that familial CCMs will almost always develop via CCM complex LOF (Figure 3 top), consistent with the lack of association between familial CCM and DVA. The probability of two CCM complex LOF mutations in trans and in the same cell (as required for CCM complex LOF in sporadic CCMs) is far lower than for a single mutation (as required for CCM complex LOF in familial CCMs). In this study we identified 6 sporadic CCMs with biallelic somatic mutations in a CCM complex gene, therefore probability of this event is likely of similar magnitude to *MAP3K3* GOF which we identified in 15 sporadic CCMs. Assuming the same relative probability for CCM complex LOF as *MAP3K3* GOF in sporadic CCMs, we determine that sporadic CCM are at least 5x more likely to develop via *PIK3CA* GOF than CCM complex LOF or *MAP3K3* GOF (Figure 3 bottom). This finding is consistent with the strong association between DVA and sporadic CCM.

**Figure 3:**
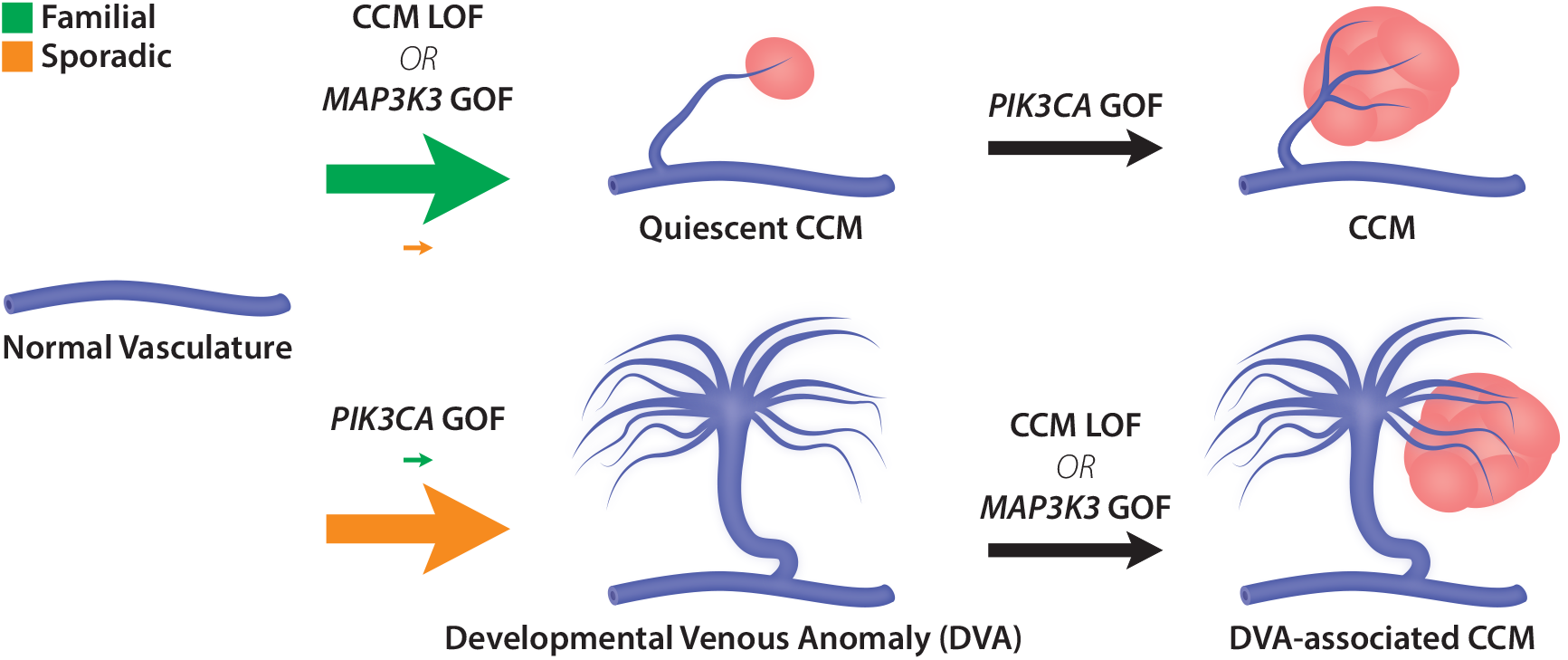
Genetic model of CCM pathogenesis. The genetic trajectories that underly familial and sporadic CCM pathogenesis. The initial path of familial and sporadic CCMs are denoted by arrow colors (green and orange respectively) with the relative probability of each path denoted by the size of the arrow. Familial CCM preferentially develop via a LOF mutation in the CCM complex (*KRIT1, CCM2, PDCD10*) to develop a quiescent CCM that may acquire an additional mutation in *PIK3CA* that drives lesion growth. Sporadic CCM are more likely to develop via a DVA caused by a *PIK3CA* mutation creating a clonal population of cells that may subsequently acquire a CCM complex LOF or *MAP3K3* GOF.

## Discussion

In this study we have further interrogated the relationship between somatic mutations in *KRIT1, CCM2, PDCD10, MAP3K3*, and *PIK3CA* which contribute to the pathogenesis of CCM. We find that somatic mutations in *MAP3K3* are not present in CCMs from individuals with familial CCM, consistent with a recent study^11^. We find that sporadic CCMs may harbor mutations in *MAP3K3, KRIT1, CCM2*, or *PDCD10*, but that the lesion will only have mutations in one of these genes. This implies that mutations in any of *MAP3K3, KRIT1, CCM2*, or *PDCD10* are sufficient for CCM formation, without the need for mutations in a second gene. As *KRIT1, CCM2*, and *PDCD10* are all members of the CCM signaling complex, this reflects the fact that the loss of any component of the complex prevents normal signal transduction. Similarly, as the CCM complex is a direct inhibitor of *MAP3K3* activity^21^, this pathway may be activated by either CCM complex LOF or by *MAP3K3* GOF, but the mutual exclusivity of mutations in these genes suggests that only one of these events is necessary for lesion formation.

CCMs often develop as the result of multiple somatic mutations that co-exist within the same cells as we show with snDNA-seq. Although several somatic mutations occur in every cell division^32^, the specificity of the mutations in CCM translates to a very low chance of acquiring these mutations within a single cell. This is especially true of somatic mutations in *MAP3K3* and *PIK3CA*, both of which have very narrow spectra of activating mutations. Despite this improbability, the accumulation of these mutations in CCM seems to occur frequently. One possible explanation for this phenomenon is that CCMs have an increased rate of somatic mutations. There is little evidence supporting this theory and such a mechanism is difficult to conceive in cases of sporadic CCM where individuals have no known genetic predisposition to CCM. An alternative explanation is that after an initial somatic mutation, the singly-mutated cell undergoes clonal expansion to form an intermediate lesion. In this study we identify 7 CCMs with either biallelic LOF in a CCM complex gene or *MAP3K3* GOF in the absence a *PIK3CA* mutation, suggesting that *PIK3CA* activation is not required for CCM formation. Furthermore, previous work in mouse models has shown that loss of a CCM complex gene (with WT *Pik3ca*) leads to clonal expansion of the mutant cells and in later stages of growth, incorporation of WT endothelial cells^33,34^. As a result of this clonal expansion, the probability of creating a double-mutant cell increases by a factor of the clonal population size as there are more cells in which the second mutation may occur. Somatic mutations in this clonal population may be acquired by genome replication during expansion or after expansion as mutations accumulate via DNA damage and error-prone repair as has been reported in other quiescent cell types^35^.

The data presented in this study suggest that DVA are an intermediate lesion as posited by the latter explanation. Genetic analysis of DVA and paired CCM show that at least some DVA develop following a somatic activating mutation in *PIK3CA*. Furthermore, plasma miRNA analysis of individuals with DVA-associated CCM revealed differentially expressed miRNAs that putatively target *PIK3CA* and *MAP3K3*, the same genes that are mutated in these lesions. By contrast, individuals with DVA but no CCM differentially expressed miRNAs that putatively target *PIK3CA*, but not *MAP3K3*. These data are consistent with the distribution of mutations we observe in CCM and DVA, and the differential regulation of these miRNAs may reflect an attempt to compensate for dysregulation of PI3K and MAPK signaling due to the somatic mutations.

The presence of *PIK3CA* mutations in DVA suggests that DVA act as a genetic precursor to CCM, which would account for the strong association between sporadic CCM and DVA (Figure 3). Likewise, DVA are not associated with familial CCM because the presence of an inherited germline mutation in a CCM gene strongly biases probability towards a CCM gene somatic mutation occurring first, as there exist many different mutations that may cause LOF, but far fewer that would cause GOF in *PIK3CA* (Figure 3B).

This study is the first to collect CCM and associated DVA for genetic analysis. Collecting tissue from CCM-associated DVA is challenging; however, collecting tissue from DVA not associated with CCM is yet more challenging as DVA are considered benign and are therefore not resected. We have attempted to address this limitation by studying biomarkers of PI3K activity which can be assayed noninvasively in blood plasma. Assaying the presence of *PIK3CA* mutations in DVA not associated with CCM will be the domain of future studies, but the data we present here demonstrate a clear link between DVA and *PIK3CA*, and suggest a model that explains the long recognized—but poorly understood—association between CCM and DVA.

While we are unable to address the presence of *PIK3CA* mutations in DVA not associated with CCM, it is worth noting that DVAs have been associated with other PI3K-related disorders^36–42^ including some cancers and neurological malformations, suggesting that DVA may have a role, possibly even as a genetic primer, in these other diseases.

## Materials and Methods

### Sample Collection

Surgically resected CCMs were obtained from the University of Chicago, the Barrow Neurological Institute, and the Angioma Alliance biobank. Additional DVA tissue was discretely dissected from the lesion during surgical resection of the associated CCM at the University of Chicago. This study was approved by each institution’s respective Institutional Review Board.

### DNA Extraction

DNA from CCM and DVA samples was extracted using the DNeasy blood and tissue kit (QIAGEN, catalog number 69504) per the manufacturers protocol. DNA purity was determined by Nanodrop and concentration was determined using the Qubit dsDNA BR assay kit (Invitrogen, catalog number Q32850) per the manufacturers protocol.

### Droplet Digital PCR

Detection of *MAP3K3* p.I441M was performed via ddPCR using a previously described probe set^20^. Assays were performed using 30-100ng of DNA with the QX200 AutoDG system (BioRad) and quantified with the QX200 droplet reader (BioRad). Analysis was performed with the QuantaSoft software (BioRad).

### Sequencing

A total of 8 sporadic CCMs with no identified mutation in *KRIT1, CCM2, PDCD10*, or *MAP3K3* (5001, 5005, 5006, 5022, 5024, 5036, 5078, and 5081) were used for whole-exome sequencing prepared using the SureSelect Human All Exon V7 probe set (Agilent, Design ID S31285117) per the manufacturers protocol. Prepared libraries were sequenced on one lane of a NovaSeq 6000 S4 flow cell for a mean depth of 133x.

### Sequence Analysis

Sequencing data was processed using the Gene Analysis Toolkit (GATK, Broad Institute) while following the GATK best practices for somatic short variant discovery using Mutect2. Secondary variant detection was performed using go-nomics (https://github.com/vertgenlab/gonomics) and bcftools mpileup to manually examine *KRIT1, CCM2, PDCD10*, and *MAP3K3* for somatic variants. Putative variants were annotated using Funcotator (GATK), the catalog of somatic mutation in cancer (COSMIC), and the genome aggregation database (gnomAD). Putative variants were filtered according to the following criteria: greater than 50x total coverage, less than 90% strand specificity, greater than 5 reads supporting the alternate allele, greater than 1% alternate allele frequency, less than 1% population allele frequency, and predicted protein/splicing change.

### Single-Nucleus DNA Sequencing

Nuclei isolation, snDNA-seq, and analysis were performed as previously described^10^.

### miRNA Extraction and Sequencing

Total plasma RNA was extracted from the plasma of 12 individuals with a sporadic CCM and an associated DVA (CCM + DVA), 6 individuals with DVA and without a CCM (DVA only), and 7 healthy controls using the miRNeasy Serum/Plasma Kit (Qiagen, Hilden, Germany) following the manufacturer isolation protocol. Diagnosis of CCM with an associated DVA, as well as DVA without a CCM lesion was confirmed on susceptibility weighted MR imaging. Illumina small RNA-Seq kits (Clontech, Mountain View, CA, USA) were then used to generate cDNA libraries, and sequencing was completed with the Illumina HiSeq 4000 platform (Illumina, San Diego, CA, USA), with single-end 50bp reads, at the University of Chicago Genomics Core. Differential miRNA analyses were completed between (1) CCM + DVA to DVA only and then (2) DVA only to healthy controls. The differentially expressed miRNAs were identified having P < 0.05, FDR-corrected. All analyses were completed using the sRNAToolbox and DESeq2 R packages^43,44^.

### Identification of Putative Targets

miRWalk 3.0 was used to identify the putative gene targets of each of the DE miRNAs, using a random forest tree algorithm with a bonding prediction probability higher than 95% on the 3 different gene locations (3’ UTR, 5’ UTR, and CDS)^45^. Putative gene targets of the DE miRNAs were identified in at least 2 of the 3 databases. DE miRNAs between (1) CCM + DVA and DVA only as well as (2) DVA only and healthy controls were then analyzed for potential targeting of the PIK3, AKT, and MAPK gene families.

### Estimating Possible LOF Mutations in KRIT1, CCM2, and PDCD10

The majority of LOF mutations in *KRIT1, CCM2*, and *PDCD10* result in either: creation of a premature stop codon (nonsense); disruption of a canonical splice site; or a frameshift. The first two of these are typically caused by a single nucleotide variant (SNV) and can be determined from the gene sequence. However, frameshift variants resulting from insertions and deletions may occur in any exon regardless of sequence context. This prevents a meaningful comparison between indel and SNV events, without making assumptions about the rate of somatic SNV and indel events. To determine a conservative lower bound for the number of LOF mutations in the CCM complex genes, we only consider nonsense and splice site mutations. The number of nonsense and splice site mutations were determined from the sequences of *KRIT1* (ENST00000394507.5), *CCM2* (ENST00000258781.11), and *PDCD10* (ENST00000392750.7). Nonsense mutations were considered any SNV in the coding sequence that would result in an in frame stop codon prior to the stop codon in the reference sequence. Splice site mutations were considered any SNV in the nearly invariant sequences at each exon-intron boundary that mediate canonical splicing.

---

**Figure S1:**
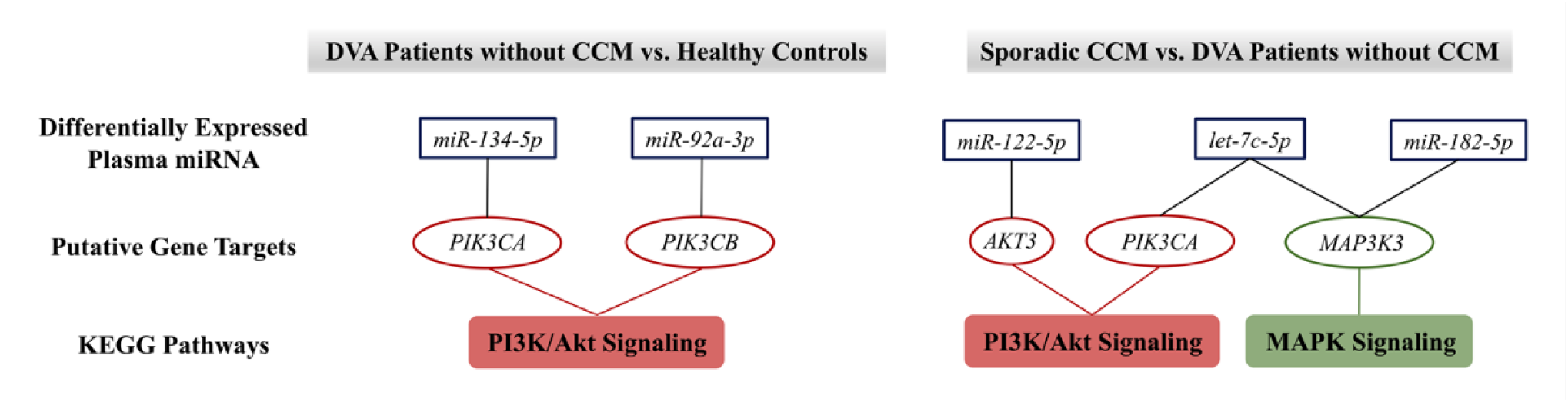
The differential plasma miRNome reflects specific pathways involved in the pathogenesis of sporadic CCMs and DVAs. Two miRNAs differentially expressed (DE) in the plasma of individuals with DVA without a CCM, *miR-134-5p* and *miR-92a-3p*, putatively target genes in the Kyoto Encyclopedia of Genes and Genomes (KEGG) PI3K/AKT signaling pathway. In the plasma of individuals with sporadic CCMs with an associated DVA, *miR-122-5p, miR-182-5p*, and *let-7c-5p* were DE and found to target genes within both PI3K/AKT and MAPK signaling KEGG pathways. All results were P < 0.05; false discovery rate corrected.

## Supplemental

### Sporadic CCM Transcriptome

Laser micro-dissected neurovascular units (NVUs) from sporadic CCM lesions were sequenced for transcriptomic analyses in comparison to micro-dissected NVUs from human brain microvasculature. The differential analyses identified 426 DE genes (DEGs) (P < 0.05; FDR corrected; FC > |1.5|). Additionally, 8 pathways from the Kyoto Encyclopedia of Genes and Genomes (KEGG) were enriched (P < 0.05, FDR corrected).

### Sporadic Transcriptome and KEGG Pathway Analyses

Six human sporadic CCM lesions were surgically resected prior to embedding in optimal cutting temperature, snap-freezing, and storing at -80°C. Control brain tissue was collected during autopsy from three subjects lacking neurological disease, fixed in formalin, and embedded in paraffin blocks. Five-µm tissue sections were mounted on Leica glass slides (Leica Biosystems Inc) and were stained, in accordance with the manufacturer’s protocols, with HistoGene (Applied Biosystems) for frozen tissue and Paradise stain (Applied Biosystems) for paraffin-embedded tissue. The neurovascular units (NVUs) from sporadic CCM lesions and normal brain capillaries were then collected using laser capture microdissection and stored at -80°C. RNA was isolated using an RNA extraction kit (RNeasy Micro Kit, Qiagen). cDNA libraries were then generated with low-input strand-specific RNA-Seq kits (Clontech) and sequenced using the Illumina HiSeq 4000 platform with single-end 50-bp reads. Differentially expressed genes were defined as P < 0.05, FDR corrected.

Enriched Kyoto Encyclopedia of Genes and Genomes (KEGG) pathways^28,29^ were obtained for the DE genes (DEGs) in the sporadic transcriptome with fold change values greater than 1.5 (P < 0.05, FDR corrected). The sporadic transcriptome was compared to the putative targets of the DE miRNAs between (1) CCM + DVA and DVA only as well as (2) DVA only and healthy controls to obtain a set of overlapping genes. Enriched Kyoto Encyclopedia of Genes and Genomes (KEGG) pathways were obtained for the overlapping genes, using a database and knowledge extraction engine with a Bayes factor greater than 3.

